# Variation in social organization and supergene control along a latitudinal gradient

**DOI:** 10.1101/2025.07.20.665811

**Authors:** Jessica Purcell, Giulia Scarparo, German Lagunas-Robles, Madison Sankovitz, Mari West, Zul Alam, Alan Brelsford

**Affiliations:** Department of Entomology, University of California, Riverside, CA, USA; Institute of Organismic and Molecular Evolution, Johannes Gutenberg Univesität, Mainz, Germany; Department of Evolution, Ecology, and Organismal Biology, University of California, Riverside, CA, USA; Department of Biology, Indiana University, Bloomington, IN, USA; Biofrontiers Institute, University of Colorado, Bounder, CO, USA

## Abstract

Widespread species often experience vastly different environmental conditions across their range. In species with polymorphic traits under strong genetic control, we can investigate how environmental variation influences the distribution of traits and whether the strength of genetic control is consistent across environmental gradients. Here, we investigate the distribution of phenotypic and genetic variation in the ant *Formica podzolica* across 30° of latitude. This species shares a supergene that is associated with colony queen number and colony sex ratio with other congeners, but we detect a surprising six common supergene variants. Colonies with a single queen are more abundant in the north compared to those with multiple queens, although the queen number polymorphism is present throughout the range. In parallel, the frequency of supergene haplotypes also varies with latitude. Of the 170 colonies examined, 10.6% contained supergene haplotypes that did not match the queen number phenotype. The highest concentration of these ‘mismatched’ colonies occurred in one site, raising the possibility that the environmental conditions there override supergene function. While the supergene system in *F. podzolica* is complex, this species holds promise for understanding how an ancient supergene system evolves in a single species experiencing highly variable environmental conditions.

## Introduction

Organismal traits frequently vary along environmental gradients, providing opportunities to study the relative contributions of phenotypic plasticity and local adaptation (reviewed by de Frenne et al. 2013, Boyle et al. 2015, Henry et al. 2023). Understanding how gene-by-environment interactions vary across a species range is empirically challenging but has been explored through modeling (e.g. Rodríguez 2013). Many empirical studies focus on investigating pairs of factors. Modern genome-wide association studies (GWAS) identify phenotype-genotype associations but often focus on phenotypic polymorphisms within one population to reduce unwanted noise derived from population structure (reviewed by Santure and Garant 2018). Landscape genomic studies seek genetic signatures of local adaptation to environmental conditions, bypassing correlated phenotypic traits (reviewed by Rellstab et al. 2015). Many ecological studies have revealed patterns in the distribution of focal traits along environmental gradients (reviewed by Purcell 2011, Halbritter et al. 2018). Here, we aim to integrate components of these fields to investigate whether and how a genotype-phenotype relationship varies across a large-scale environmental gradient.

Phenotypic polymorphisms controlled by supergenes (regions of the genome containing two or more functional mutations that shape a complex phenotype) present a novel opportunity to explore the interaction between phenotype, genotype, and environment. Recently discovered supergenes control complex polymorphic traits in species that are geographically widespread, including migration strategies in fish (e.g. Matschiner et al. 2022, Thompson et al. 2020), color and life history strategies in birds (e.g. Funk et al. 2021, Maney and Küpper 2022), and colony social organization in ants (e.g. Purcell et al. 2014, Brelsford et al. 2020, Wang et al. 2013, Yan et al. 2020). Supergenes have often been detected using GWAS (e.g. Purcell et al. 2014, Gutiérrez-Valencia et al. 2022, Jay et al. 2022, Voelker et al. 2023). In such cases, further research is needed to investigate how supergene structure and control of the focal trait vary across the species range. Inversions associated with local adaptation, some of which act as supergenes (e.g. Merót et al. 2018), have been discovered through investigations of genomic features that vary in frequency across populations (e.g. Lowry and Willis 2010, Kapun et al. 2016, Harringmeyer and Hoekstra 2022, Meyer et al. 2024, Vangestel et al. 2024). In this case, identifying clear genotype-phenotype associations can be challenging, as other genetic markers and multiple phenotypes are likely to co-vary across populations (e.g. Kapun et al. 2016, Harringmeyer and Hoekstra 2022). Combining the genotype-phenotype relationships from the GWAS approach with the broader sampling effort in the geographic survey approach provides an opportunity to investigate supergene-by-environment interactions, thereby addressing knowledge gaps remaining in both bodies of literature.

Arthropod social traits, such as colony size, frequently vary along environmental gradients (Purcell 2011), suggesting that the benefits of different social strategies depend upon their ecological context. In *Anelosimus* social spiders, for example, colony sizes are larger in the lowland tropical rainforest compared to mid- and high-elevation habitats in the Andes, both within the social species *A. eximius* (Purcell and Avilés 2007) and in a comparison of congeneric species (Avilés et al. 2007). Spiders in the lowland tropical rainforest have access to and capture larger prey, and the larger social groups are also more resilient to disturbance by predators or intense rains (Guevara and Avilés 2015). Identifying patterns in the distribution of social traits enables researchers to investigate whether these patterns derive from behavioral plasticity (e.g. Kapheim et al. 2020, Omufwoko et al. 2023), local adaptation (e.g. Purcell et al. 2015, Charmantier et al. 2016), or an interaction between the two. Even in traits with well-established supergene control, researchers have observed exceptions to genotype-phenotype relationships (e.g. McGuire et al. 2022, Scarparo et al. 2023, Scarparo et al. 2024, Lajmi et al. 2024) that may result from varying environmental conditions.

Many species within the ant genus *Formica* are socially polymorphic, with some colonies containing a single queen (=monogyne) and others containing multiple queens (=polygyne) (e.g. Sundström et al. 2005, Chapuisat 2023). As in other socially polymorphic ant systems, several individual- and colony-level traits consistently differ between monogyne and polygyne colonies (Rosset & Chapuisat 2007). A chromosome-spanning supergene, first discovered in *Formica selysi* (Purcell et al. 2014), is widespread across the *Formica* genus (Brelsford et al. 2020, Purcell et al. 2021). Individuals from polygyne colonies generally have at least one copy of a rearranged ‘P’ haplotype, while individuals from monogyne colonies generally have only the ancestral ‘M’ haplotype. Detailed studies of additional species (*Formica glacialis*, Lagunas-Robles et al. 2021; *Formica francoeuri,* Pierce et al. 2022; *Formica neoclara*, McGuire et al. 2022; *Formica cinerea*, Scarparo et al. 2023; *Formica aserva,* Scarparo et al. 2024), revealed that each species exhibits notable differences in the distribution of genotypes within colonies. Nevertheless, we expect all socially polymorphic *Formica* species to have total or partial supergene control of colony queen number.

Across the range of a widely distributed species, the extent of supergene control may vary depending on the local environment if the costs and benefits of alternative strategies differ in different habitats. In the *Formica* supergene system, reproductive strategies are known to differ between monogyne and polygyne colonies (e.g. Fontcuberta et al. 2021). Specifically, new queens of monogyne origin are more likely to participate in a mating flight and then establish a nest independently, while queens of polygyne origin are more likely to mate near their natal nest and join established colonies. If one strategy is favored in a given environment, this could lead to local adaptation in the form of selection favoring one supergene haplotype over the other.

Alternatively, a plastic response to unpredictable environmental conditions could override supergene control in some cases. For example, if queens of monogyne origin colonize an environment with few available nest sites and high queen mortality, newly mated queens that join existing nests may have better fitness outcomes than those that attempt to disperse and establish new nests. Along the same lines, queens of polygyne origin may be more successful at independent nest founding in habitats with abundant resources and nest site availability, even if they have fewer fat reserves than queens of monogyne origin. In proximate terms, local conditions may influence the expression of certain genes within the supergene, facilitating a more plastic response to the environment. If the strength of supergene-trait associations varies across different environments, this phenomenon would be overlooked by relying on supergene genotype as a proxy for social structure (or any other trait of interest).

In this study, we take advantage of the broad geographic range (Francoeur 1973) and known social polymorphism (Deslippe & Savolainen 1995, Deheer & Herbers 2004) of the North American species *Formica podzolica* to investigate how environmental variation along a latitudinal gradient shapes phenotypic and genotypic polymorphism. We sequence samples from sites spanning 30° of latitude to ask whether the phenotype (the frequency of monogyny and polygyny), the genotype (the frequency and diversity of supergene haplotypes), and the genetic control of phenotype vary consistently with latitude. We predict that monogyny will be more prevalent at higher latitudes, consistent with an earlier study showing that monogyny increases in frequency at higher elevations in *F. selysi* (Purcell et al. 2015). With such an extensive geographic range, we expect some variation in the frequency of supergene haplotypes across different environmental conditions. We do not have an *a priori* prediction of whether and how genetic control will vary across the species range. Since the latitudinal gradient is a proxy for numerous correlated environmental gradients, we additionally investigate whether observed patterns are correlated with temperature or precipitation. Finally, we ask whether and how underlying population structure might contribute to supergene and phenotype distribution patterns.

## Methods

### Focal species

*Formica podzolica* is a widespread North American ant species found in boreal forests, often associated with aspen groves. The species builds conspicuous clay or dirt mounds ranging from 25 cm to several meters in diameter. Previous studies in Alberta (Canada) and Colorado (USA) demonstrated that the species exhibits a social polymorphism in queen number, with populations containing a mix of monogyne and polygyne colonies (Deslippe & Savolainen 1995, Deheer & Herbers 2004). More recently, Lagunas-Robles et al. (2021). discovered that the colony-level production of males or future queens in a *F. podzolica* population from Yukon Territory, Canada is associated with variation in a supergene. Previous studies have not established whether the variation in colony queen number is also controlled by a supergene in this species.

### Field sampling

Members of the Purcell and Brelsford labs collected *F. podzolica* workers for this study from 2015-2022. We conducted two systematic sampling trips that together spanned 30+ degrees of latitude, from the southwestern United States to the Arctic Circle in Alaska (2016: California to Alaska; 2021: Arizona and New Mexico to Montana). These trips focused on sampling *Formica* diversity throughout the western United States and Canada, and we collected ants in two complementary ways. First, we visually searched for colonies and collected 8-20 workers from the nest entrance or on the dome of the nest. Second, we collected single workers along transects, where we sampled the first *Formica* worker encountered and then the next worker encountered at least 100 m away along a road or trail. We sampled eight ants per transect. We collected additional samples from colonies during projects focused on other aspects of *F. podzolica* behavioral ecology in Alberta (West and Purcell 2020), Colorado (Sankovitz and Purcell 2021), and Utah, Colorado, and Alberta (Lagunas-Robles, unpublished data).

### DNA extraction

We extracted genomic DNA from the head and thorax of workers or the head of alates (gynes or males) using Qiagen DNeasy blood and tissue kits, following the insect tissue protocol (until 2021) or the QiaAmp 96 kit with a QiaCube HT extractor robot (since 2021). In both cases, we made some minor protocol adjustments to reduce cost and/or increase yield. In both cases, we cut the portion of the ant to be extracted using a flame-sterilized scalpel and then thoroughly ground the tissue with a pestle after immersing the tissue (within a 1.7 mL tube) in liquid nitrogen. We retained the unextracted portion of each ant as a voucher in 100% ethanol. For both methods, we incubated the tissue in a Proteinase K solution overnight before proceeding to the DNA wash and elution steps. For the DNeasy kit, we used alternatively sourced spin columns (BPI-tech), completed the second DNA wash with 70% ethanol, and eluted in 30 μL of EB buffer. For the QiaAmp kit, we followed the manufacturer’s protocol but eluted in 100 μL of EB buffer.

### Whole genome sequencing

We sequenced the genomes of *F. podzolica* workers (N = 278), males (N = 20), and gynes (N = 2) collected from states and provinces from New Mexico to Alaska (sampling sites and sequencing details summarized in Table S1). We completed the library preparation using Seqwell Plexwell LP-384 kits, following the manufacturer’s protocol. We sequenced pooled libraries containing *F. podzolica* and other non-focal *Formica* species on an Illumina Novaseq 6000 at Novogene Inc. or on an Illumina Novaseq 6000 or an Illumina HiSeq X Ten at the UCSD Institute for Genomic Medicine. We removed the non-focal species from downstream analyses.

### RAD sequencing

We collected restriction-site-associated DNA (RAD) sequence data on 1054 *F. podzolica* workers sampled across the species range using the protocol of Brelsford et al. (2016). In brief, we digested the genomic DNA of each individual using two restriction enzymes (Table S2), ligated inline barcoded adapters to one restriction site and universal adapters to the other, and used Sera-Mag magnetic beads (Millipore Sigma) in a PEG/NaCl buffer (Rohland and Reich 2012) or Mag-Bind magnetic bead solution (Omega Bio-tek) to remove small DNA fragments. We then amplified restriction fragments in four replicate PCR reactions for each individual, adding a plate-specific index, and ran a final extension step with additional dNTP and primers to reduce the frequency of single-stranded PCR products. Before pooling, we ran the product on an agarose gel to identify failed samples. We then pooled successfully amplified samples from each plate and ran a final magnetic bead cleanup to remove small DNA fragments. We pooled libraries (up to 28 plates in one pool) containing *F. podzolica* and other non-focal *Formica* species and sent them to genomic core facilities for sequencing. In this case, we sequenced *F. podzolica* samples in five different sequencing batches (batch information is summarized in Table S2). We removed non-focal species from downstream analyses.

### Whole genome data preparation

We removed adapter sequences from raw reads and merged overlapping paired-end reads using *PEAR* v0.9.11 (Zhang et al. 2014). We aligned raw reads to the newly assembled chromosome-level reference genome of *F. glacialis*, a close relative of *F. podzolica* (see “*Formica glacialis* genome assembly” section below for assembly methods), using *Bwa-mem2* v2.2.1 (Vasimuddin et al. 2019), and removed PCR-duplicate reads with *Samtools* v1.18 (Li et al. 2009; Danecek et al. 2021). We used *Bcftools* v1.18 (Li 2011; Danecek et al. 2021) to call variants, ignoring reads with mapping quality < 20. We then filtered the raw variant calls with *VCFtools* (Danecek et al. 2011), retaining genotypes with read depth of at least 1 and single nucleotide polymorphisms (SNPs) with genotype calls in at least 80% of individuals and a minor allele count of at least 3. We excluded indel variants, SNPs with more than two alleles, and SNPs with heterozygosity greater than 75%, resulting in a filtered dataset of 4,554,091 SNPs.

We called variants in the mitochondrial genome for all *F. podzolica* individuals in the WGS dataset using *Bcftools* mpileup (Li 2011; Danecek et al. 2021), specifying a haploid variant calling model, ignoring reads with mapping quality < 40, retaining invariant sites, and adding two previously published *F. glacialis* and one *F. subsericea* individual genomes (*F. glacialis* NCBI accessions SRR15979804 and SRR15979820, *F. subsericea* NCBI accession SRR15979803; Purcell et al. 2021) as an outgroup. We removed individuals with high rates of missing data. We retained loci with genotype quality of at least 20 in at least 80% of remaining individuals, producing a data matrix with 7,710 positions in 284 individuals.

### RADseq data preparation

We merged paired reads using *PEAR* v0.9.11 (Zhang et al. 2014) and aligned reads to the chromosome-level *F. glacialis* genome using *Bwa-mem2* v2.2.1 (Vasimuddin et al. 2019). We then used *Samtools* v 1.18 (Li et al. 2009; Danecek et al. 2021) to sort and index the resulting bam files and called variants using *Bcftools* mpileup (Li 2011; Danecek et al. 2021) for each sequencing batch, ignoring reads with mapping quality < 20 for most batches and with a minimum mapping quality of < 40 for one batch (‘ger’) that included samples with more variable DNA yield. We filtered the raw variant calls within each batch independently with *VCFtools* (Danecek et al. 2011). Specifically, we applied a minimum depth filter of 7, retained SNPs with genotype calls in at least 80% of individuals, and those that occurred at a minor allele frequency of at least 5%. We also excluded indels, removed SNPs with more than 2 alleles, and excluded SNPs with heterozygosity greater than 75%.

### Identification of supergene haplotype variation

Using the whole genome sequencing data, we employed two complementary approaches to identify supergene variants. First, using *Plink* 1.9 (Purcell et al. 2007), we ran principal component analyses (PCAs) on variable markers from two sections of the supergene: the previously identified M_D_ region (Lagunas-Robles et al. 2021; 2.05-5.37 Mbp in the *F. glacialis* genome assembly, see genome assembly section below for details) and the remaining portion of the supergene region originally identified in *F. selysi* (14.06-19.09 Mbp in the *F. glacialis* genome assembly). Second, we inspected variable sites in a 15 kb region around the gene “*Knockout”*, which was previously found to contain haplotype-specific SNPs across many different *Formica* species (Brelsford et al. 2020; Purcell et al. 2021).

We calculated pairwise F_ST_ in 10-kbp windows between individuals from each set of PCA clusters (*i.e.*, individuals that were in the same clusters in both PCA analyses and had similar variation around *Knockout*). We used the empirical boundaries around regions of high F_ST_ to calculate PCAs within each region for each RADseq batch. We assessed RADseq batches separately both because we used different enzyme combinations for several batches (Table S2) and because we have previously encountered moderate batch effects (e.g. Scarparo et al. 2023). We also identified SNPs that were diagnostic between haplotypes, thereby confirming genotype calls from the RAD-based PCAs. This dual approach enabled us to confidently call genotypes for all sequenced individuals.

### Colony social structure

To infer the social structure of colonies, we used *CoAncestry* (Wang 2011) with Wang estimator to calculate nestmate pairwise relatedness values in all colony-based collections with at least five individuals sequenced with RADs (N = 170 colonies). We calculated pairwise relatedness values for all individuals within each batch to avoid an upward bias in nestmate relatedness values due to a batch effect, except for the ‘ger’ batch (Table S2), which was further subdivided by locality (Alberta, Colorado, Utah) to minimize bias from population structure. In all cases, nestmates were all sequenced in the same batch. We removed all markers on chromosome 3 to ensure that queen number inferences were independent of the supergene. We visualized pairwise relatedness values for each colony (Figure S1) and assigned colonies as monogyne monandrous when all individuals were related at a level of 0.5 or above, as monogyne polyandrous when at least 40% of workers were highly related (0.5 or higher) and none had relatedness below 0.2, and as polygyne when at least two individuals had relatedness below 0.2 or when the proportion of highly related individuals (0.5 or higher) was below 40%.

### Climatic data collection

We downloaded WorldClim data (Fick and Hijmans 2017) at the 30s scale and used the raster command in R (Package raster, Hijmans and van Etten 2012) to attach values for WorldClim variables 1-19 to our collection sites for both the whole genome and RADseq datasets. For each dataset, we ran a PCA using the prcomp command in R to determine which variables had the greatest weighting on PC axes 1 and 2. We visualized the vector weightings with the fviz_pca_var command in R (Package factoextra, Kassambara and Mundt 2020). Across the two PCAs, PC axis 1 was generally aligned with temperature measurements, while PC axis 2 aligned with precipitation measurements. In both datasets, bio4 (temperature seasonality) and bio16 (precipitation during the wettest quarter) were closely aligned with PC axes 1 and 2, respectively. Thus, we used these variables to assess how temperature and precipitation levels influence the distribution of colony social structure and supergene variants.

### Population structure

We investigated population structure across the *F. podzolica* range by running *Admixture* v1.3.0 (Alexander et al. 2009) on our whole genome sequencing dataset with K values from 2 to 10. We excluded all SNPs on chromosome 3 and pruned SNPs on other chromosomes based on linkage disequilibrium following the recommendations in the *Admixture* user manual, using the Plink flag “--indep-pairwise 50 kb 10 0.1”, resulting in a dataset of 579,844 SNPs. Cross-validation error was lowest for K=2, separating northern from southern samples. We estimated Weir and Cockerham’s F_ST_ between these two populations using *Vcftools* (Danecek 2011) and identified 440 SNPs with F_ST_ > 0.9.

We inferred a maximum-likelihood tree of mitochondrial genomes using *IQ-TREE* v2.2.2.6 (Minh et al. 2020), with the TPM3u+F+I+R3 model chosen by the *ModelFinder* component of the pipeline (Kalyaanamoorthy et al. 2017).

### Statistical analysis

We used logistic regression models, implemented in R 4.1.3 (R Core Team 2022) with the glm function using binomial error structure, to test whether colony social structure, with monogyny coded as 0 and polygyny coded as 1, varies consistently with latitude and WorldClim variables bio4 (temperature seasonality) and bio16 (precipitation of the wettest quarter). We used multinomial logistic regression, implemented in R with the multinom function in the nnet package (v 7.3-17, Venables and Ripley 2002), to test the association between supergene haplotypes and latitude, bio4, and bio16. We similarly used glm models with binomial error structure to test whether the presence of any P haplotype (P_L_, P_S_, and P_NM_), the P_L_ only, or the P_S_ only, respectively, predicted whether a colony was monogyne or polygyne. In these models, we included latitude along with presence of any P, P_L_ only, or P_S_ only as fixed effects.

### Formica glacialis genome assembly

We assembled a chromosome-level reference genome for *F. glacialis*, a close relative of *F. podzolica*, using PacBio and Omni-C sequence data and a linkage map. We sent multiple flash-frozen males from monogyne colony BJFC6 in Big John Flat, Utah to Dovetail Genomics for DNA extraction and sequencing. Dovetail extracted high-molecular-weight DNA from a pool of males, obtained 33.2 Gbp of PacBio circular consensus sequence data, and produced an initial assembly of the PacBio data using *Hifiasm* v0.15.4-r347 (Cheng et al. 2021) with default parameters. Dovetail identified non-arthropod contigs in the assembly with *Blobtools* v1.1.1 (Laetsch et al. 2017) and removed them. They then identified and removed duplicate contigs representing alternative copies of the same genomic region with *purge_dups* v1.2.5 (Guan et al. 2020).

Dovetail produced an Omni-C proximity ligation library from male ants from the same colony and sequenced the library to approximately 30x depth on an Illumina Hiseq X. They then used the *HiRise* software package (Putnam et al. 2016) to scaffold the initial PacBio assembly.

To obtain independent information about large-scale genome structure in *F. glacialis*, we constructed a high-density linkage map. We collected 62 workers from monogyne colony SYLC5 in Sylvan Lake, Alberta, extracted DNA, and produced RAD sequence data using the procedures described above, with restriction enzymes PstI and MseI. We aligned the resulting reads to Dovetail’s HiRise genome assembly using *Bwa-mem* v0.7.17 (Vasimuddin et al. 2019) and called variants using *Bcftools* v1.15 (Li 2011; Danecek et al. 2021). We filtered variants using *Vcftools* v0.1.17 (Danecek et al. 2011), retaining genotype calls with phred-scaled genotype quality > 20 and SNPs with genotype calls retained in at least 80% of individuals and minor allele frequency > 0.1. We then identified SNPs heterozygous in >80% of genotyped individuals using a custom awk script and removed them using *VCFtools*. We reformatted the data for *MSTmap* (Wu et al. 2008), including each SNP twice with opposite phases to account for the unknown phase of the mapping family (Gadau 2009). We then constructed a linkage map using *MSTmap* with a significance threshold of 1e-5 and default values for other parameters.

Comparing the linkage map and HiRise assembly, we noted several cases where distinct linkage groups had been merged in the HiRise scaffolding process and two cases of intrachromosomal rearrangement (see Results). Because the number of linkage groups (27) matched the expected number of chromosomes in non-parasitic *Formica* species (Hauschteck-Jungen and Jungen 1976), we interpreted these mismatches as errors in the HiRise scaffolding process. To correct the scaffolding errors, we manually split the improperly joined contigs at the location of the nearest gap and produced new chromosome-level scaffolds that matched the linkage map. We aligned at least ten segments of each corrected scaffold to the published *F. selysi* reference genome (Brelsford et al. 2020) using *Bwa-mem2* and named the *F. glacialis* scaffolds based on their homologous *F. selysi* scaffolds. We visualized synteny between the HiRise assembly, linkage map, and final re-scaffolded assembly with *NGenomeSyn* (He et al. 2023).

To assemble the *F. glacialis* mitochondrial genome, we used 25 bp segments of mitochondrial COI and CytB sequence from the *F. podzolica* mitochondrial genome (Allio et al. 2020) to query *F. glacialis* raw PacBio reads. We extracted five reads starting near COI and ending near CytB (a distance of approximately 9 kbp of the expected 16 kbp circular genome), and five reads from the same strand starting near CytB and ending near COI (completing the coverage of the circular genome). We manually aligned these ten reads and obtained the majority-rules consensus for each nucleotide in the alignment.

We assessed genome completeness using conserved hymenopteran single-copy orthologs in *BUSCO* v5.8 (Simao et al. 2015, Waterhouse et al. 2018, Seppey et al. 2019, Manni et al. 2021).

### Genome annotation and identifying genes of interest

We transferred gene annotations produced by the Global Ant Genome Alliance (Vizueta et al. 2025) for the published reference genome of *F. selysi* to the new *F. glacialis* assembly using *LiftOff* (Shumate and Salzberg 2021) with default parameters. We then used *bedtools* (Quinlan and Hall 2010) to obtain lists of genes within supergene regions of interest and those within 5 kbp of any SNP that was highly differentiated (F_ST_ > 0.9) between the northern and southern genetic clusters.

## Results

### Formica podzolica has six widespread supergene haplotypes

Based on whole genome data, we detected six distinct supergene haplotypes on chromosome 3 (Figure 1). Overall, there were two ‘M’ haplotypes, the ‘sex ratio supergene’ M_A_ and M_D_ described by Lagunas-Robles et al. (2021). There were four ‘P’ haplotypes that differ in their extent across chromosome three (Table 1).

**Figure 1:**
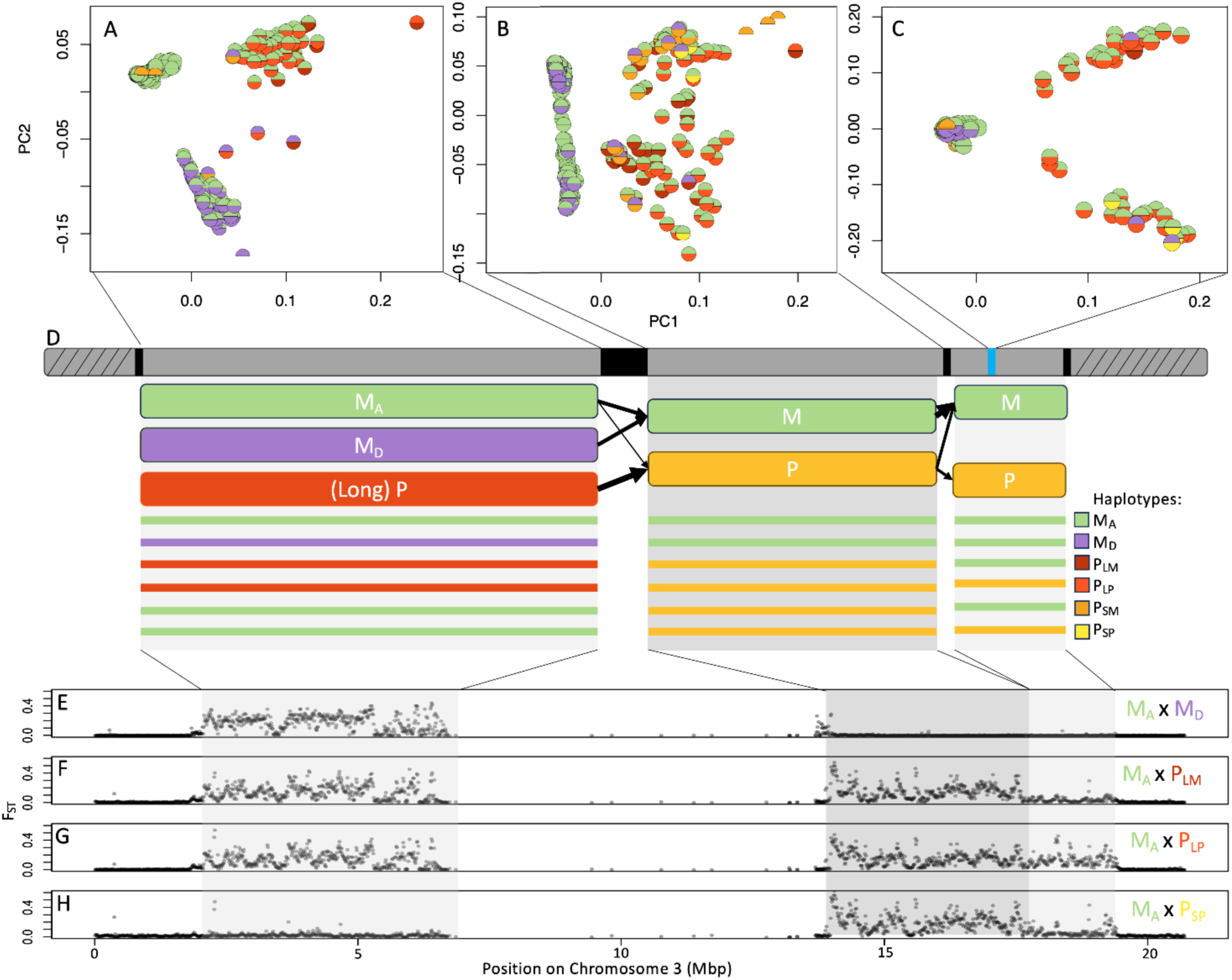
In the whole genome dataset, we identified six distinct supergene haplotypes on chromosome 3. These haplotypes differ from one another across three distinct cassettes of chromosome 3. PCAs of markers in each cassette (A, B, and C spanning three regions denoted in panel D) separate individuals with M_A_, M_D_, and P haplotypes in the cassette left of the centromere (A), individuals with M and P haplotypes in the cassette just to the right of the centromere (B), and individuals that have the M and P versions of the cassette spanning the gene *Knockout* (C), shown as a blue line in panel D. In the PCAs, the two haplotypes of each individual are color-coded within each half-circle. The haplotype compositions are illustrated in the diagram showing each of the three regions of differentiation (D). Genetic differentiation (F_ST_) along chromosome 3 between M_A_M_A_ homozygotes and individuals heterozygous for M_A_ and: M_D_ (E), P_LM_ (F), P_LP_ (G), and P_SP_ (H) is displayed in the lower panels. The central region of chromosome 3, from ∼6-14 Mbp, contains a large region of centromeric repeats.

**Table 1:**
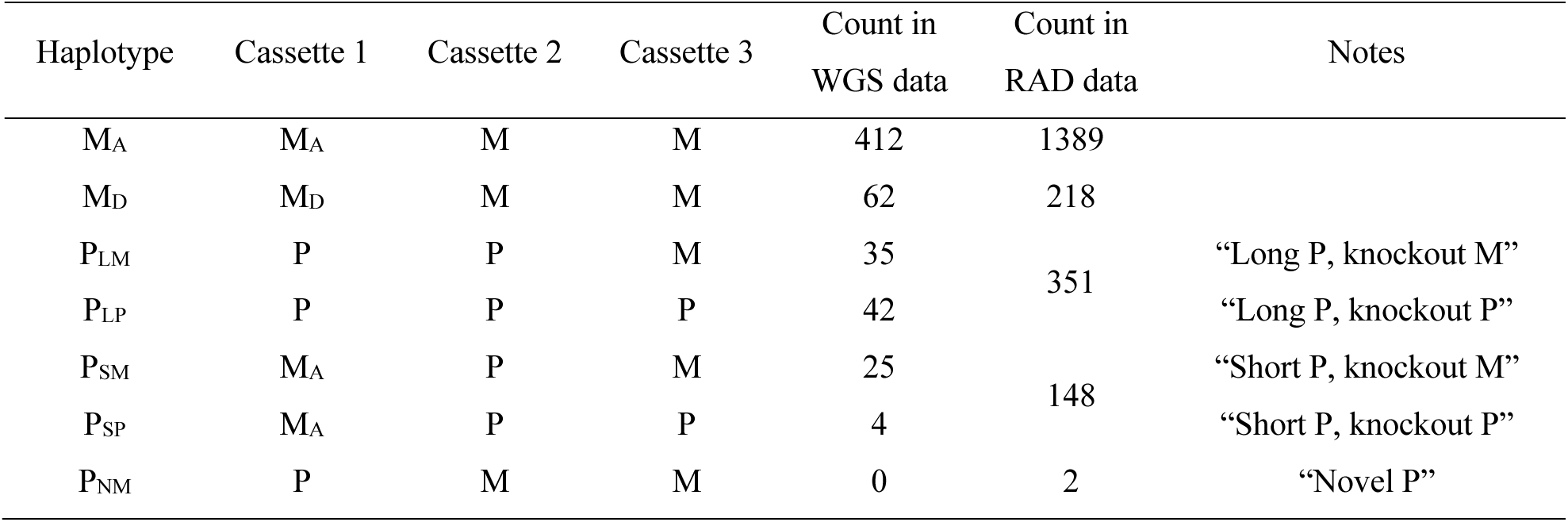
Haplotype names and compositions, as well as sample sizes within the whole genome (WGS) and RADseq datasets. Note that cassette 3 is small and ‘knockout M’ versus ‘knockout P’ could not be reliably distinguished in the RADseq libraries, so counts for P_LM_ + P_LP_ and P_SM_ + P_SP_ are combined in the ‘Count in RAD data’ column.

The six supergene haplotypes spanned three major regions of variation, or “cassettes”. The first cassette, based on the M_D_ region described by Lagunas-Robles et al. (2021), extends from ∼2 to 6 Mbp and exhibits three variants, M_A_, M_D_, and P (Figure 1A, Table 1). In one PCA cluster, individuals had excess homozygosity (M_A_/M_A_), while individuals in the other two clusters had excess heterozygosity in this region (M_A_/M_D_ and M_A_/P, Table 2). The genome assembly revealed a putative centromere spanning chromosome 3 from ∼6-14 Mbp, where short reads do not align accurately due to the repetitive nature of the satellite DNA sequence. The second cassette, spanning from ∼14 to 17.5 Mbp, exhibited two different variants, M and P, separated along PC1 (Figure 1B). Intriguingly, some individuals that were M_A_/M_A_ or M_A_/M_D_ in the first cassette appeared to be M/P (with excess heterozygosity) in the second cassette. This result revealed two alternative putative P haplotypes: one spanning the ∼2-17.5 Mbp region, which we called “long P” (P_L_), and one spanning the 14-17.5 Mbp region, which we called “short P” (P_S_). Our inspection of a small region (15 kb) around the gene *Knockout* revealed a third cassette with two variants (separated along PC1, Figure 1C), and substantial differences associated with broader population structure in one of the variants (separated along PC2, Figure 1C). The latter variant was only found in individuals that were heterozygous in the second cassette (M/P), while the former was found in both individuals that were homozygous (M/M) and heterozygous (M/P) in the second cassette (Figure 1C). This was an unexpected finding, because our assessment of other species revealed consistent M- and P-associated *Knockout* variants that shared 22 haplotype-diagnostic SNPs among highly divergent species (Brelsford et al. 2020; Purcell et al. 2021). While M/M *F. podzolica* individuals always had the ‘M’ variant in the *Knockout* cassette, both P_L_ and P_S_ could either have the ‘M’ or the ‘P’ variant. We distinguished these by adding either an ‘M’ or a ‘P’ to the subscript (Table 1). For both cassettes depicted in Figures 1B and 1C, PC2 revealed within-haplotype variation, distinguishing individuals collected in northern and southern populations. The names and compositions of each haplotype are summarized in a schematic diagram (Figure 1D, see also Table 1), with differentiation between genotypes shown in a series of pairwise Fst comparisons (Figure 1E-H).

**Table 2:**
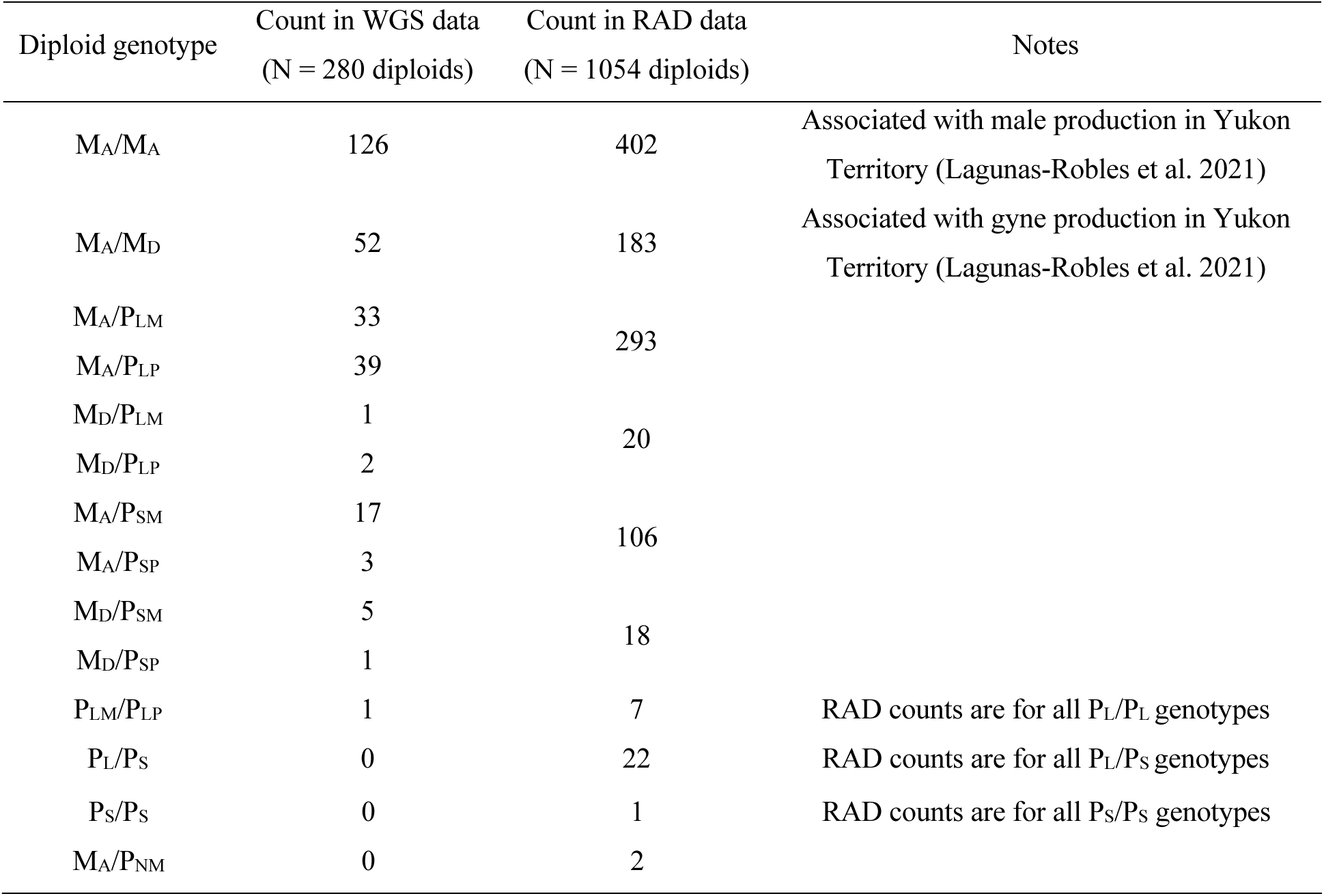
Genotypes and sample sizes within the whole genome (WGS) and RADseq datasets (for diploid individuals only). Note that cassette 3 is small and ‘knockout M’ versus ‘knockout P’ could not be reliably distinguished in the RADseq libraries, so counts for P_LM_ + P_LP_ and P_SM_ + P_SP_ are combined in the ‘Count in RAD data’ column.

### Geographic variation in haplotype distribution

Although most supergene variants are widely distributed, they differed significantly in their latitudinal distribution (Figure 2A). We combined the P_LM_ and P_LP_ haplotypes into one category, P_L_, and combined the P_SM_ and P_SP_ haplotypes into a second category, P_S_, for statistical analyses due to small sample sizes. The P_L_ haplotype was found at significantly lower latitudes than the other haplotypes (Figure 2A, compared to M_A_, p = 0.0035; compared to M_D_, p = 0.0004; compared to P_S_, p = 0.0030; other pairwise comparisons were not significantly different). While our sample was not large enough to conduct a quantitative assessment, we noted that we never found the rare P_SP_ variant north of 45° N. Latitude, *per se*, is unlikely to influence genotype distributions, so we investigated correlated abiotic variables accessed from WorldClim. A PCA on samples used for whole genome sequencing (Figure 2B) showed that sampling localities for *F. podzolica* vary in temperature (axis 1) and precipitation (axis 2), with temperature seasonality (bio4) and precipitation rate in the wettest quarter (bio16) each strongly associated with PC axis 1 and 2, respectively. Individuals with a P_L_ haplotype tended to occur in less seasonally variable environments than M_D_ and P_S_ haplotypes (Figure 2C, compared to M_D_, p = 0.0077; compared to P_S_, p = 0.0088) and in habitats with lower precipitation rates in the wettest quarter (Figure 2D, compared to M_A_, p = 0.0115; compared to M_D_, p = 0.0033; compared to P_S_, p = 0.0003). P_S_ tended to occur in habitats with higher precipitation rates in the wettest quarter compared to M_A_ (p = 0.0180).

**Figure 2.**
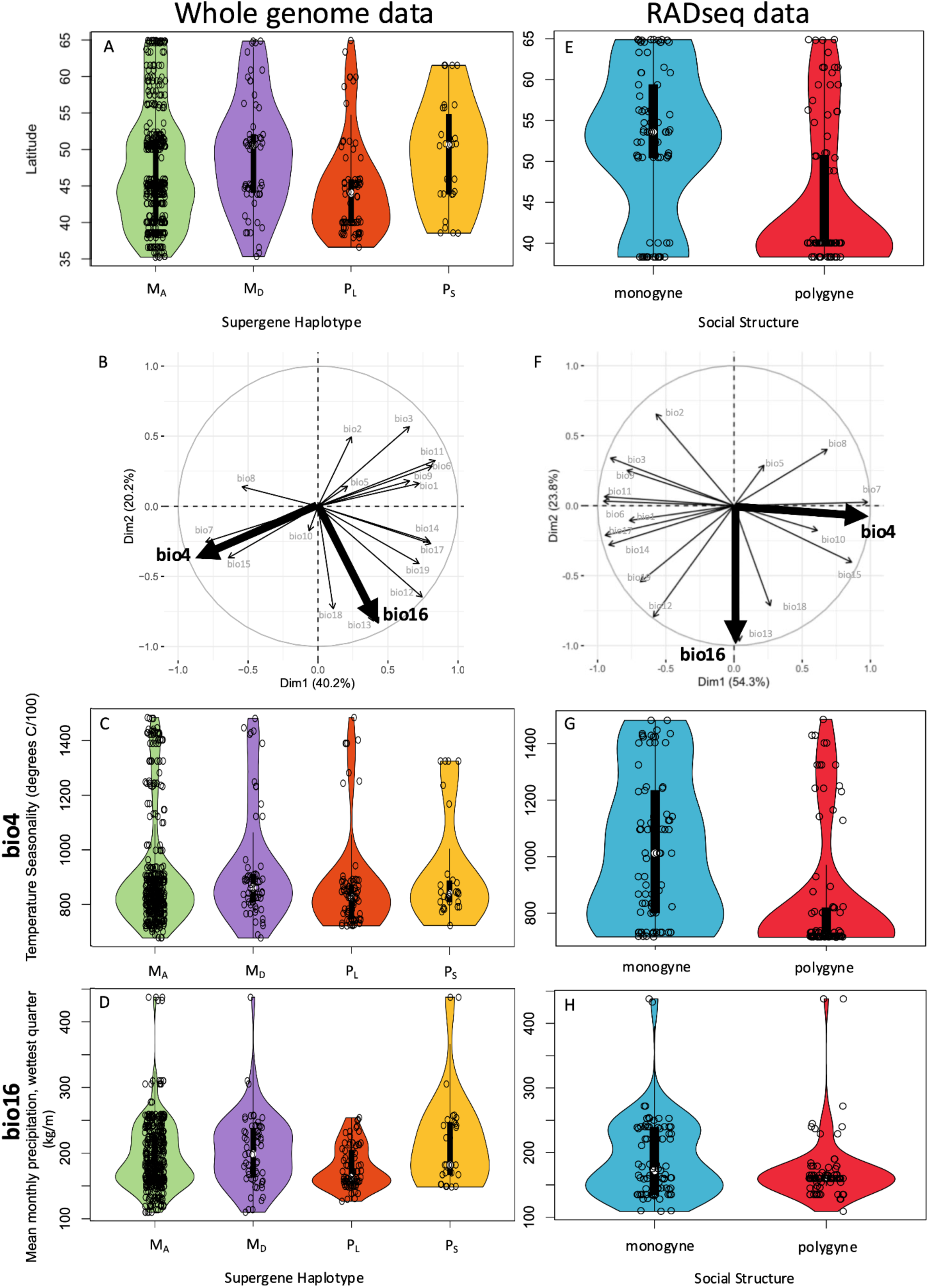
The distribution of supergene haplotypes (based on whole genome sequencing, left panel) varies significantly with latitude, with the P_L_ haplotype distributed significantly further south on average than the other three haplotypes (A). A PCA of WorldClim data (B) reveals that temperature and precipitation across the *F. podzolica* habitat are orthogonal. The P_L_ haplotype also occurred in less seasonally variable environments compared to the M_D_ and P_S_ haplotypes (C) and in drier environments (during the wet quarter) compared to all three haplotypes (D). The P_S_ occurred in wetter environments than the M_A_ haplotype. Social structure also varies significantly along the latitudinal gradient, with monogyne colonies being more common in northern latitudes (E). The PCA of WorldClim based on the RADseq data (F) resembles the results based on whole genome data (B). Social structure varies more strongly with temperature (G) than precipitation (H).

### Monogyny is more prevalent at northern latitudes

The latitudinal distribution of monogyne (including both monandrous and polyandrous colonies) and polygyne colonies differed significantly (Figure 2E). Pairwise relatedness values among nestmates from 170 colonies distributed from Utah and Colorado in the south to Alaska and Yukon Territory in the north revealed a pattern of polygyny being more common in the south and monogyny being more common in the north (Figure 2E; logistic regression: z = -4.785, df = 167, p < 0.0001).

We investigated abiotic factors associated with the distribution of social organization. As with the whole genome dataset (Figure 2B), sampling localities for *F. podzolica* used in the RADseq analyses varied in temperature (axis 1) and precipitation (axis 2) in a PCA (Figure 2F). We assessed the social structure distribution with respect to temperature seasonality (bio4) and precipitation rate in the wettest quarter (bio16). Monogyne colonies tended to occur in more seasonally variable environments (Figure 2G, logistic regression: z = -4.388, df = 167, p < 0.0001) and in areas with greater precipitation rates during the wettest quarter (Figure 2H, logistic regression: z = -2.017, df = 167, p = 0.044).

### Despite range wide patterns, phenotypic and genetic polymorphism are widespread

In almost all well-sampled sites (with at least five colonies analyzed), the *F. podzolica* population was socially polymorphic, with only one site in central Alberta (N = 8 colonies) containing exclusively monogyne colonies. Likewise, almost all sites contained both M and P haplotypes (Figure 3).

**Figure 3.**
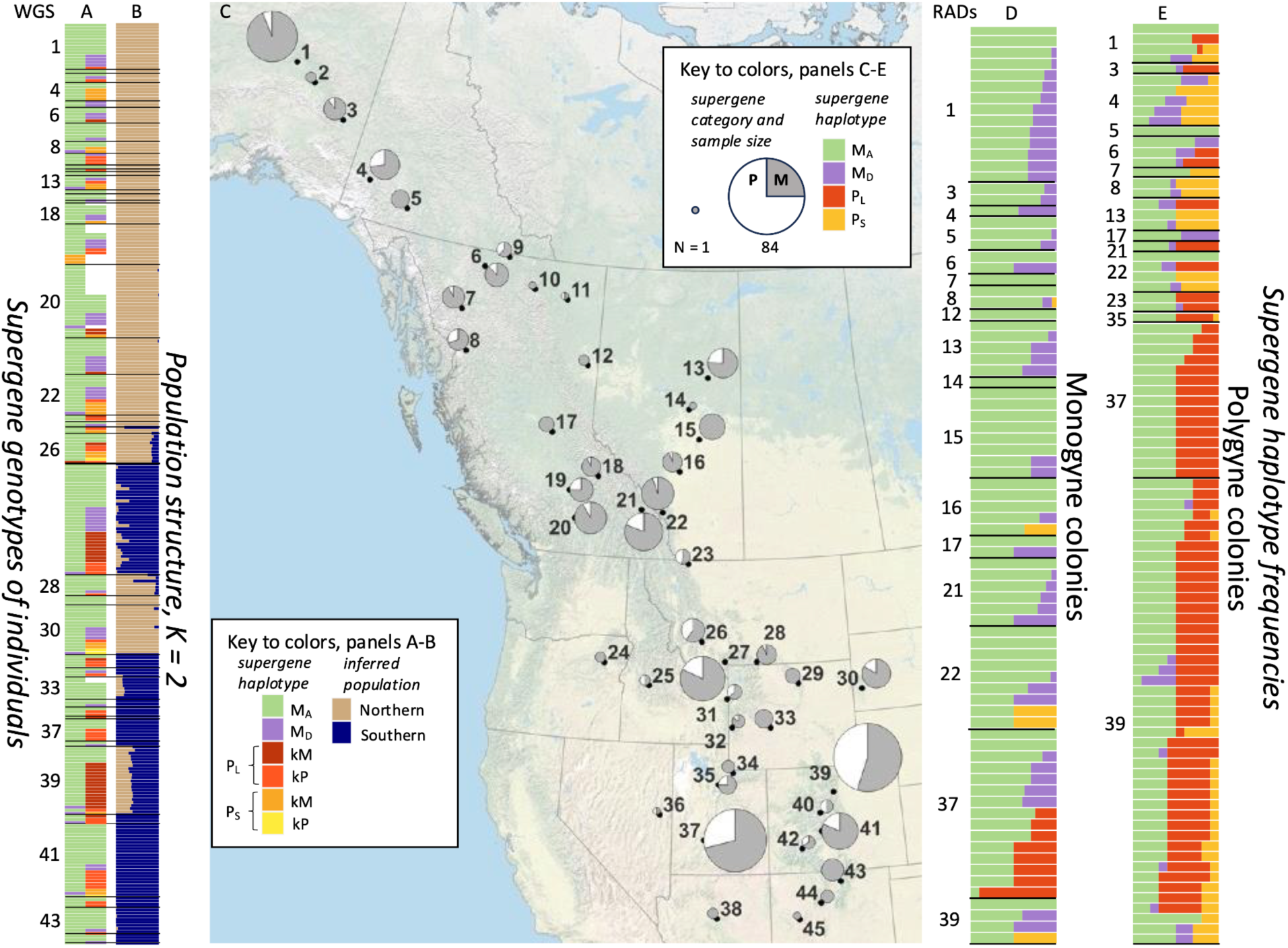
Supergene genotypes (A) and inferred population membership (based on K = 2 in *Admixture*, B) of individuals (each bar in the two plots shows one individual) reveal that the majority of individuals are either homozygous M_A_M_A_ or heterozygous for M_A_ and one of the other haplotypes (A). P_S_ and M_D_ variants are more common in the north, while P_L_ is more common in the south. The population is structured along a north-south axis (B), with admixture observed in northern Colorado (site 39), southern Montana (26-28), and central Wyoming (33). The map (C) shows sampling effort from Alaska (sites 1-3) in the north to New Mexico (43-45) in the south and combines both whole genome and RADseq datasets. Pie charts are scaled by sample size and show the distribution of M (M_A_ + M_D_) and P (P_L_ + P_S_) haplotypes, revealing that most sampling localities have multiple supergene variants, even in the northern part of the range where both polygyny and the P haplotype are relatively rare. Bar charts to the right of the map show the proportion of supergene haplotypes within colonies (each bar represents a colony with at least five workers genotyped) determined to be monogyne (D) or polygyne (E). The association of M haplotypes with monogyny (D) and P haplotypes with polygyny (E) is clear despite some exceptions. Notably, many apparently monogyne colonies in Utah (site 37) contain one or more individuals with a P_L_ haplotype, while five polygyne colonies from northern populations lack any P haplotypes.

**Figure 4.**
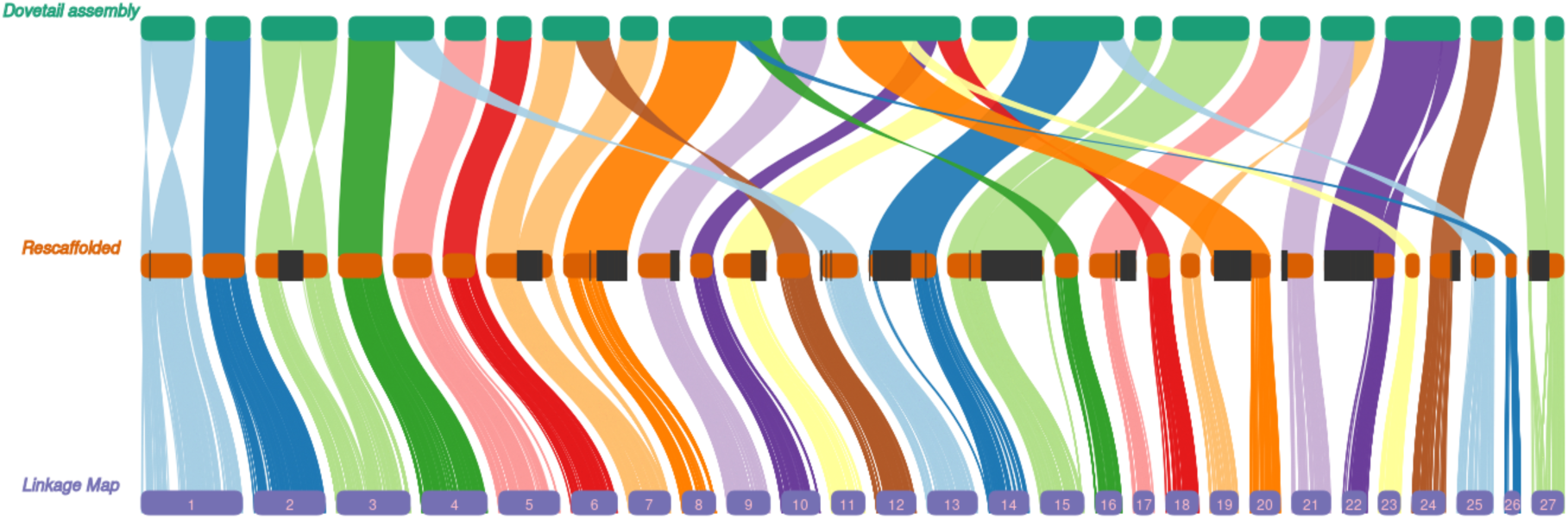
The *F. glacialis* genome assembly produced by Dovetail Genomics includes some large-scale assembly errors. Errors included spurious fusions, fissions, and inversions, often located at or near arrays of centromere-associated satellite DNA (shown with grey bars in the central panel). Rescaffolding the genome using a high-density linkage map resulted in a marked improvement in the chromosome-level assembly.

With six common haplotypes present in the WGS dataset, there could be as many as 21 distinct genotype combinations in diploids. We found ten genotypes in the WGS dataset, with all haplotypes except M_A_ occurring mainly in a heterozygous state (Figure 3A, Table 2). Additional genotypes and one additional haplotype were detected in the RADseq dataset, but these were rare (Tables 1, 2).

### Genotype x Phenotype x Environment interactions

Overall, the presence of a P haplotype (whether P_L_ or P_S_) is significantly correlated with polygyny (presence of any P: z = 6.63, df = 167, p = 3.3*10^-11^, latitude NS; presence of P_L_: z = 6.12, df = 167, p = 9.5*10^-10^, latitude NS; presence of P_S_: z=4.89, df = 167, p = 1.0*10^-6^, latitude z = -4.46, p=8.2*10^-6^). Thirteen of the 81 colonies inferred to be monogyne contained at least one individual with a P haplotype, while five of the 89 colonies inferred to be polygyne contained no individuals with a P haplotype. Among the 13 P haplotype-containing monogyne colonies, eight were found at a single site in central Utah, had the P_LP_ haplotype, and comprised more than half of the monogyne colonies detected at the site. The remaining five were distributed across the species range and had other P haplotypes. In contrast, the five polygyne colonies lacking a P haplotype were distributed in Canada and Alaska. Even within polygyne colonies where the P haplotype is present, some individuals were homozygous M_A_/M_A_; these colonies existed throughout the range.

### Genetic population structure

The *F. podzolica* population exhibits the most pronounced genetic differentiation between northern and southern populations (*Admixture* K = 2, Figure 3B). Sites in Montana, Wyoming, and northern Colorado (sites 26-28, 33, and 39, Figure 3C, Table 3) were admixed between the northern and southern populations.

**Table 3:**
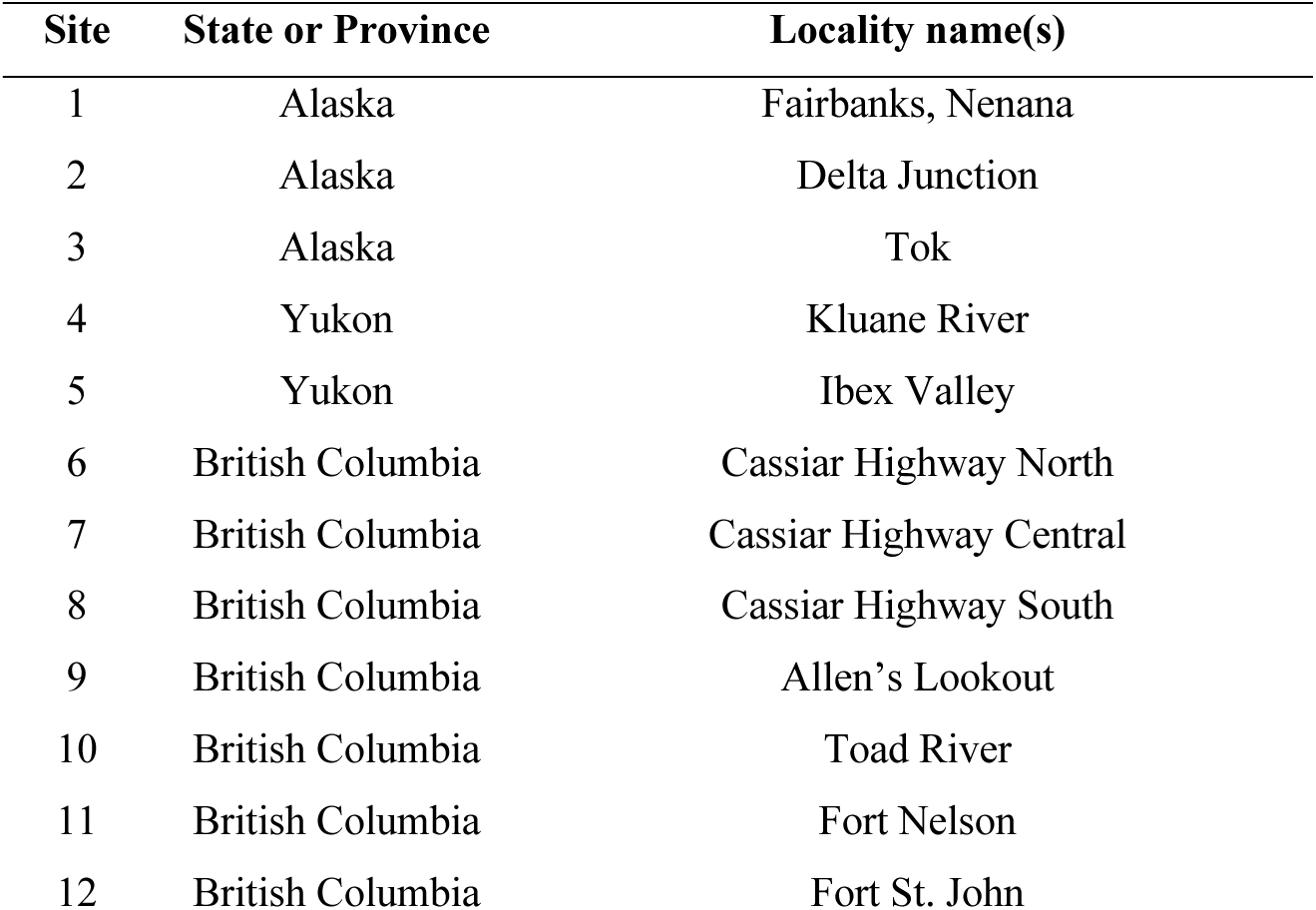
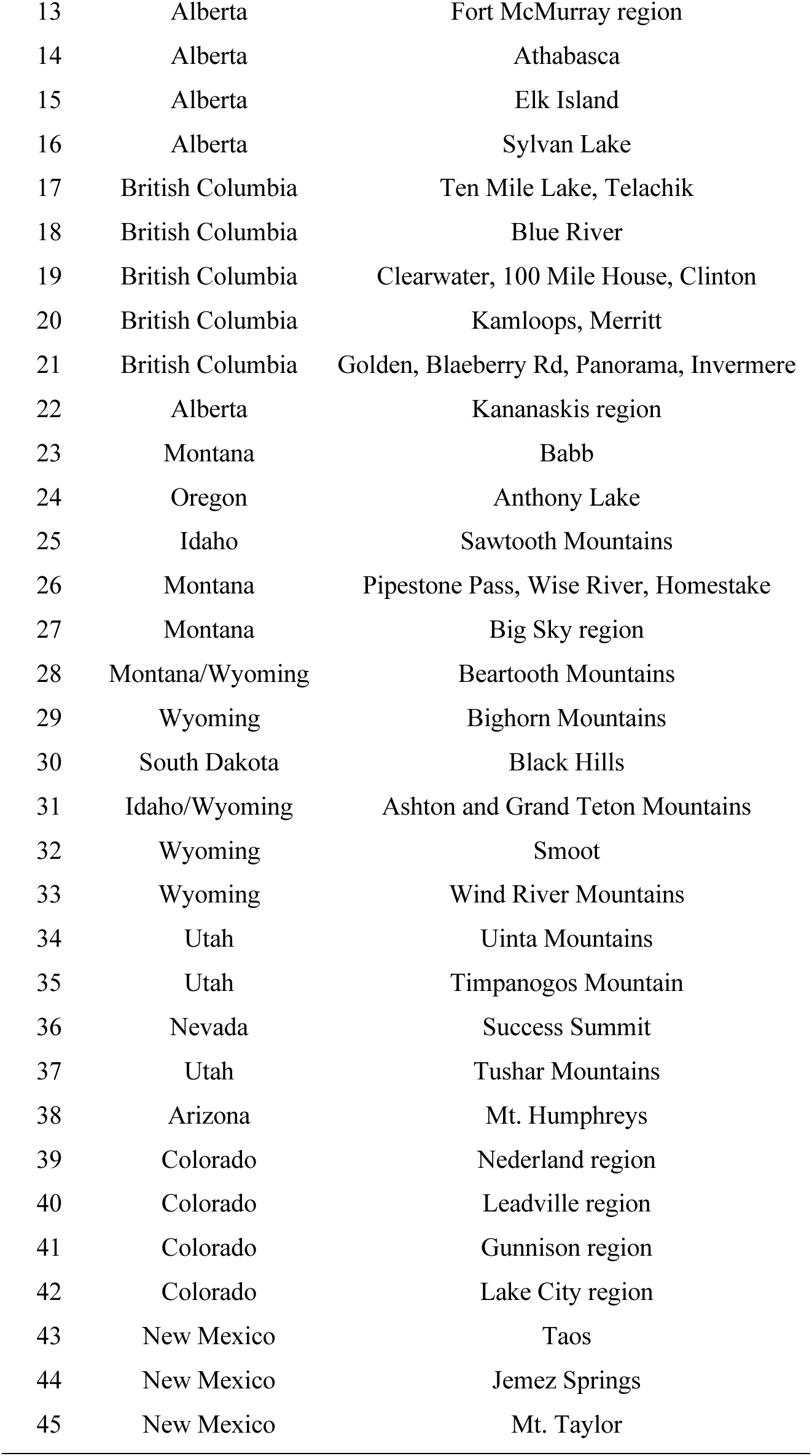
List of site numbers and locality names; note that site numbers match those shown on the map in Figure 3C.

Patterns of population structure based on the mitochondrial genome revealed some contrasting patterns with the nuclear genome-based analysis (Figure S2). As in the nuclear analysis, northern and southern populations showed the greatest mitochondrial differentiation. Within the southern mitochondrial clade, there was clear substructure separating populations from southern Colorado and Utah (sites 35-37, 40-42). Interestingly, samples from Taos, NM (site 43) include a mitochondrial haplotype found only in Taos and several haplotypes similar to those found in southern Colorado and Utah. A second clade containing individuals with southern and mixed nuclear ancestry was found in Idaho, western Wyoming, and parts of Montana (sites 26-28, 31-33). Within the northern mitochondrial clade, one subclade contained only samples from the Black Hills in South Dakota (site 30), while the remaining subclade included samples from Canada and Alaska (sites 1-22) as well as the isolated population from eastern Oregon (site 24).

### Genome assembly

The *F. glacialis* genome assembly was 469.7 Mbp in length, with 27 chromosomal scaffolds, 118 unplaced scaffolds, and one mitochondrial contig. Of the total assembly length, 142.3 Mbp (30.3%) was on the unplaced scaffolds, which were largely comprised of a 129 bp satellite motif associated with centromeres in other *Formica* species (Lorite et al. 2004). The assembly N50 was 11.1 Mbp. Our genome assembly had an overall BUSCO completeness score of 96.5%.

### Genes in supergene cassettes

The entire supergene region, spanning chromosome 3 from 2 to 19.5 Mbp, contained a total of 487 genes in the annotated *F. glacialis* genome. The first cassette (M_D_ region) contained 200 genes (Table S3), the second cassette contained 172 genes (Table S4), and the third cassette (*knockout* region) contained 115 genes (Table S5). We note that many genes in the supergene likely do not contribute to the functional differences among haplotypes. In contrast with findings in *Solenopsis invicta* (Pracana et al. 2017), we detected no odorant receptors (ORs) in the *Formica* supergene.

### Differentiated genes across the species range

We identified a total of 77 genes within 5 kbp of SNPs that were highly differentiated between the northern and southern populations (F_ST_ > 0.9, Table S6).

## Discussion

The supergene region of *F. podzolica* includes six widespread variants. Some haplotypes differ in their distribution with respect to latitude, although most overlap in one or more populations.

The social polymorphism in colony queen number is also distributed throughout the species range, but monogyne colonies are more prevalent at northern latitudes. The combination of supergene variation, a widespread phenotypic polymorphism, and gradients in the distribution of both genotypes and phenotypes sets the stage for an investigation of the interactions between genotype, phenotype, and environmental variation. The supergene variation observed in this study also offers a new perspective on the variable evolutionary trajectories of different supergene regions, providing additional context for the discussion of genotype and phenotype variation along environmental gradients.

### Supergene variants differ from one another in discrete sections of chromosome 3

We detected six relatively widespread supergene haplotypes (plus an additional rare haplotype found only in northern Colorado), which is the most intraspecific variation yet described in an ant supergene system (Lagunas-Robles et al. 2021 described three haplotypes; Scarparo et al. 2023 described four haplotypes). These haplotypes consist of distinct combinations of three different cassettes along chromosome 3 (Figure 1D). For example, the differences between the M_A_ and M_D_ haplotypes were found in the first cassette (from ∼2 to 6 Mbp), as previously described by Lagunas-Robles et al. (2021). Differences between the P_L_ and P_S_ haplotypes similarly mapped to the first cassette, with P_L_ differentiated from both M haplotypes, but P_S_ appearing indistinguishable from M_A_ in that region. In the second cassette (from ∼14-17.5 Mbp), both P haplotypes (P_L_ and P_S_) and both M haplotypes (M_A_ and M_D_) cluster together, but the M and P clusters are strongly differentiated from each other. We suggest that some of these alternative haplotypes have formed through rare double recombination (or gene conversion) events homogenizing parts of the supergene region during the evolutionary history of *F. podzolica*. This hypothesis is consistent with the ‘eroding strata’ model of supergene evolution proposed by Brelsford et al. (2020) to explain why few regions of an apparently ancient supergene contain SNPs that consistently distinguish M and P haplotypes across divergent *Formica* species.

The third cassette contains the gene *Knockout*, which was previously identified as a candidate gene for shaping the queen number polymorphism, as 22 SNPs consistently distinguished individuals of monogyne and polygyne origin in five polymorphic species (Brelsford et al. 2020). Given the previous findings, we were surprised to discover that both P_L_ and P_S_ variants in *F. podzolica* were associated with either ‘*Knockout P’* or ‘*Knockout M’*. Moreover, all P haplotypes (with either version of *Knockout*) were significantly associated with polygyny. This association suggests that *Knockout* is not an essential gene in the development or maintenance of monogyny and polygyny, at least in *F. podzolica*. Moreover, the ‘modular’ nature of the supergene variation will serve as a natural experiment to investigate which genes in each region of the supergene shape components of the polygyny syndrome. While we do not have the resolution to investigate these questions in this study, it is possible that components of the polygyny syndrome, such as queen fecundity, body size, or dispersal strategy, vary across the four alternative ‘P’ haplotypes discovered in this system.

### Phenotype and genotype vary with latitude

Monogyne colonies are more common at northern latitudes and tend to occur in habitats with more seasonal temperature variation than polygyne colonies. Nevertheless, both social forms are present across the 30° latitudinal range of *F. podzolica.* This pattern mirrors the variation in colony social structure along elevation gradients with widespread polymorphism found in *F. selysi* (Purcell et al. 2015), albeit on a much larger geographic scale. This distribution could reflect environmentally-mediated variation in the costs and benefits of queen number or associated traits, or it could have formed as a consequence of differential colonization histories in northern latitudes by individuals of monogyne and polygyne origin following the last glacial maximum. In general, monogyne queens are expected to be more proficient dispersers (Sundström 1995; Sundström et al. 2005; Fontcuberta et al. 2021), which is consistent with a more rapid northward spread of monogyne populations.

Across the latitudinal range, the four distinct P haplotypes also differ in frequency, with the P_L_ variants generally occurring further south than the P_S_ variants. As proposed in a theoretical study by Jay et al. (2024), supergene variation across the range may be a signature of local adaptation; in this case, alternative P variants may contain locally adapted alleles encoding as yet undetermined traits. Alternatively, the distribution of P haplotypes could be an outcome of historical processes, such as an ancient introgression of a P haplotype from another species into a *F. podzolica* population or differential selection on the supergene in different glacial refugia. Since it primarily occurs in regions where polygyny is relatively rare, the P_S_ variant was less common in our samples than the P_L_. Both P haplotypes are found together in a limited number of sites, including in northern Colorado (site 39), the Black Hills, South Dakota (site 30), and Fairbanks, Alaska (site 1).

### Does the environment influence genotype/phenotype mismatches?

Overall, 10.6% of colonies show a mismatch between genotype and phenotype, with 13 monogyne colonies containing one or more individuals with a P haplotype and five polygyne colonies with no P haplotypes detected. Despite these mismatches, the presence of either the P_L_ or P_S_ haplotype is significantly associated with polygyny in this species. While there is no strong pattern of mismatch frequency varying along latitudinal gradients, environmental factors may influence supergene control of social structure in some contexts. First, most monogyne colonies containing a P haplotype are found at one locality in Utah (site 37). This site notably contained many colonies under stones, whereas *F. podzolica* usually builds clay dome nests, sometimes overlaid with small pebbles. We hypothesize that the availability of ‘safe’ nesting sites under rocks may enable more queens, including those with a P haplotype, to establish colonies independently. In addition, our team may be more likely to find incipient colonies if they are present under stones. These colonies may acquire additional queens in subsequent years. It is also worth noting that senescing polygyne colonies may appear monogyne based on an assessment of worker relatedness when they are in decline to the point of having just one reproductive queen. Across the full dataset, monogyne colonies containing P haplotypes could have either P_L_ or P_S_ variants.

The five polygyne colonies without any P-bearing individuals were found in Canada and Alaska, raising the possibility that this mismatch is favorable under some conditions at northern latitudes. For example, if nesting sites are limited or local competition is intense, monogyne groups may be more inclined to accept new queens. Interestingly, polygyne colonies containing a mix of individuals with and without P haplotypes were relatively common, a pattern also described in *F. neoclara* (McGuire et al. 2022). The samples for the five nests lacking P haplotypes included 7-9 individuals, so we think it unlikely but not impossible that we simply did not collect P-bearing individuals from these colonies. Although mismatches between genotypes within colonies and social structure are uncommon, the frequency of this pattern varies among sites in our sample. Thus, we caution researchers about inferring phenotype directly from supergene genotype in previously unsampled localities, and we suggest that these mismatches provide an opportunity to study the genetic mechanisms underlying complex phenotypes.

### Population structure

Within both the nuclear and mitochondrial genome analyses, we detect strong differentiation between northern and southern populations of *F. podzolica* (Figures 3, S2, with contact zones and nuclear admixture identified in northern Colorado (population 39) and to the west and south of Yellowstone National Park (populations 27, 31-33)). These two genetic populations may reflect past allopatry during previous glacial periods. We note that the P_L_ haplotypes are more common in the southern population and the P_S_ haplotypes are more common in the northern population, which is consistent with the possibility that the evolution of P haplotype diversity in *F. podzolica* was mediated by a history of allopatry between populations in different glacial refugia. The mitochondrial tree reveals three major clades: one containing the southernmost populations (in Colorado, Utah, Nevada, Arizona, and New Mexico), one containing most sites in the contact zone between the northern and southern populations (in Wyoming, Idaho, and central Montana) and one including individuals from the northern sites, along with the Black Hills and Bighorn mountains (Figure S2). The northern populations exhibit far less mitochondrial genetic structure than the southern populations, consistent with a relatively recent range expansion northward. A second, non-mutually exclusive hypothesis that could explain the deeper differentiation between southern populations is that these tend to occur at higher elevations, with valleys of unsuitable habitat separating them, while the forest habitat of *F. podzolica* is more continuous in the northern part of the range.

### The genes within the supergene

Among the 487 genes within the supergene region based on the *F. glacialis* genome annotation, we found at least five gustatory receptors, at least one gene involved in olfaction in *Drosophila melanogaster*, and one gene involved in juvenile hormone biosynthesis. These classes of genes have been implicated in previous studies examining the genetic basis of eusocial behaviors (e.g. Kocher & Kingwell 2024; Zhou et al. 2015). We also note that numerous genes encoding proteins involved in transcription are present in the supergene region. Additional work is needed to identify which genes in the region influence queen number and associated phenotypes, but the composition of the *F. podzolica* supergene, with three distinct cassettes across chromosome 3, could help to isolate genes found in different portions of the supergene region for functional testing.

### Conclusions

Overall, we discovered that the geographically widespread species *F. podzolica* exhibits extensive variation in the *Formica* supergene that is associated with colony queen number (Purcell et al. 2014) and colony sex ratio (Lagunas-Robles et al. 2021). Combinations of three cassettes form six common haplotypes and an additional rare haplotype. Notably, one transition between cassettes occurs at the centromere, which was detected in the newly assembled *F. glacialis* genome. Both the supergene haplotypes and the colony queen number phenotype vary in frequency with latitude, with monogyne colonies more common at northern latitudes and the P_L_ haplotypes more common at southern latitudes. Importantly, the supergene is usually but not always a good predictor of social structure; one population in Utah had a particularly high number of deviations from this genotype-phenotype association. The differential distribution of the P_L_ and P_S_ haplotypes roughly mirrors the deep population structure between northern and southern samples, which hybridize in northern Colorado and the area west of Yellowstone National Park. This unusually complicated supergene system holds promise for future research examining deviations from genotype-phenotype associations and how each portion of the supergene region influences the suite of traits related to colony queen number.

## Data Availability

Genetic sequences and the *F. glacialis* genome assembly will be available on NCBI under BioProject PRJNA1293783. Additional sample metadata can be found in the supplementary material files.

## Acknowledgements

This research was supported by the US National Science Foundation (DEB grant #1942252 to J.P. and DEB grant #1754834 to A.B. and J.P.), as well as Alberta Conservation Association Grants in Biodiversity (to M.W. and to G.L.-R.). G.L.-R. received additional funding from the Irwin M. Newell Graduate Research Fund, the Vaughn H. Shoemaker Graduate Fellowship, and a fellowship from the US National Science Foundation (DGE-1326120). M.W. and M.S. received fellowship support from the US National Science Foundation (grant #1631776), and M.S. also received support from a Boulder County Open Spaces grant. Computational work was carried out on the UCR HPCC, which has been funded by grants from the US National Science Foundation (MRI-2215705 and MRI-1429826) and the National Institute of Health (1S10OD016290-01A1). This study contains data generated at the UC Berkeley QB3 Genomics Laboratory using the Illumina HiSeq 4000, supported by National Institute of Health S10 OD018174 Instrumentation Grant, and at the UC San Diego IGM Genomics Center using the Illumina NovaSeq 6000, purchased with funding from National Institute of Health S10 OD026929. The authors thank Elk Island National Park (AB) for permission to collect, under permit # EI-2021-38658. The authors also thank Dr. Junxia Zhang for her assistance with field collection.

