## Supplementary Materials figures and text for "Variation in social organization and supergene control along a latitudinal gradient"

Supplementary Materials for: Purcell et al. Variation in social organization and supergene control along a latitudinal gradient

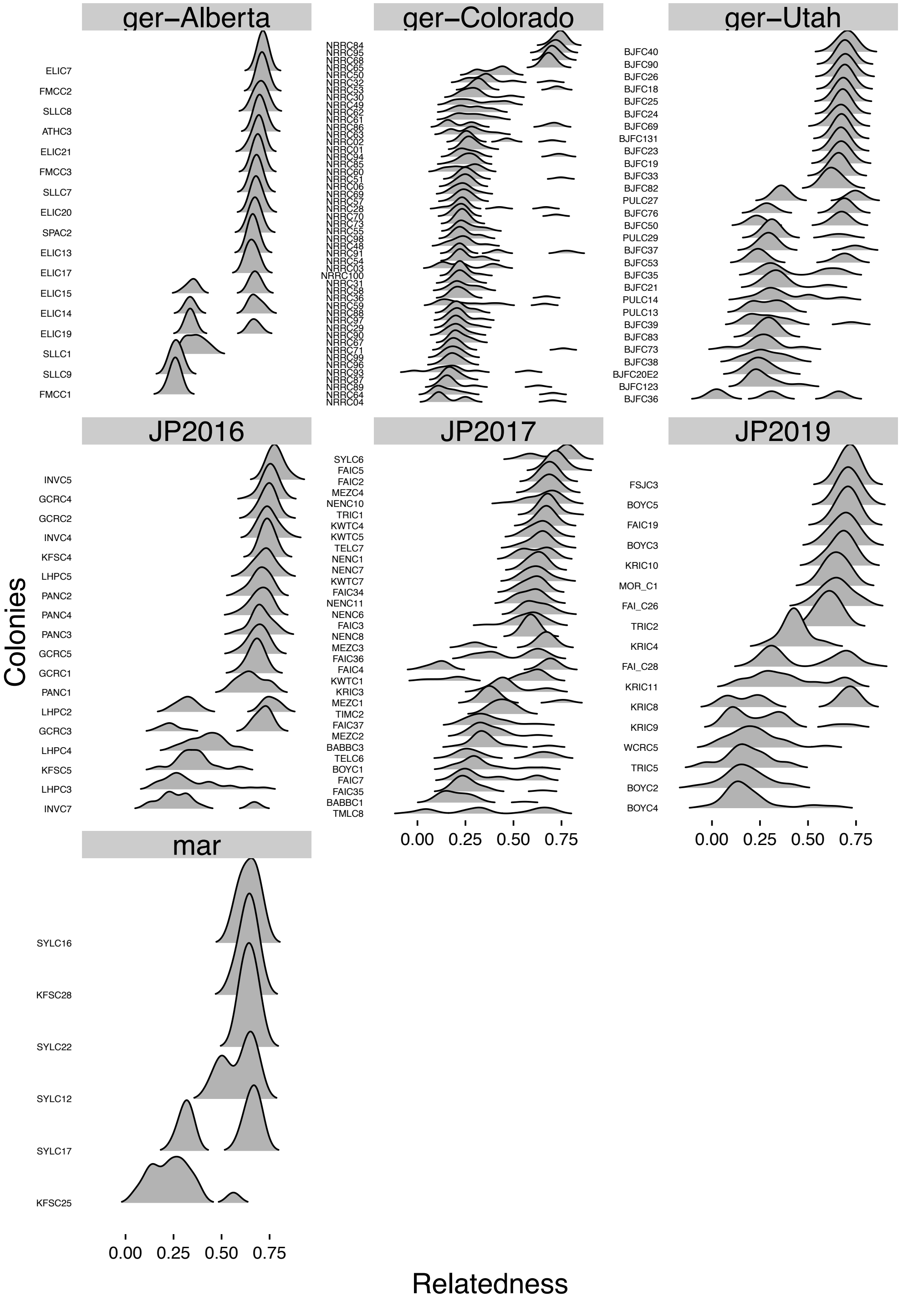


**Figure S1**: Ridgeline plots of relatedness based on RADseq data for 170 *F. podzolica* colonies. Each sequencing batch (and locality within the large *'ger'* batch) was analyzed separately. Chromosome 3 was excluded from all analyses to ensure that phenotype assessment was independent of supergene genotype. As described in the methods, we assigned colonies as monogyne monandrous when all individuals were related at a level of 0.5 or above, as monogyne polyandrous when at least 40% of workers were highly related (0.5 or higher) and none had relatedness below 0.2, and as polygyne when at least two individuals had relatedness below 0.2 or when the proportion of highly related individuals (0.5 or higher) was below 40%.

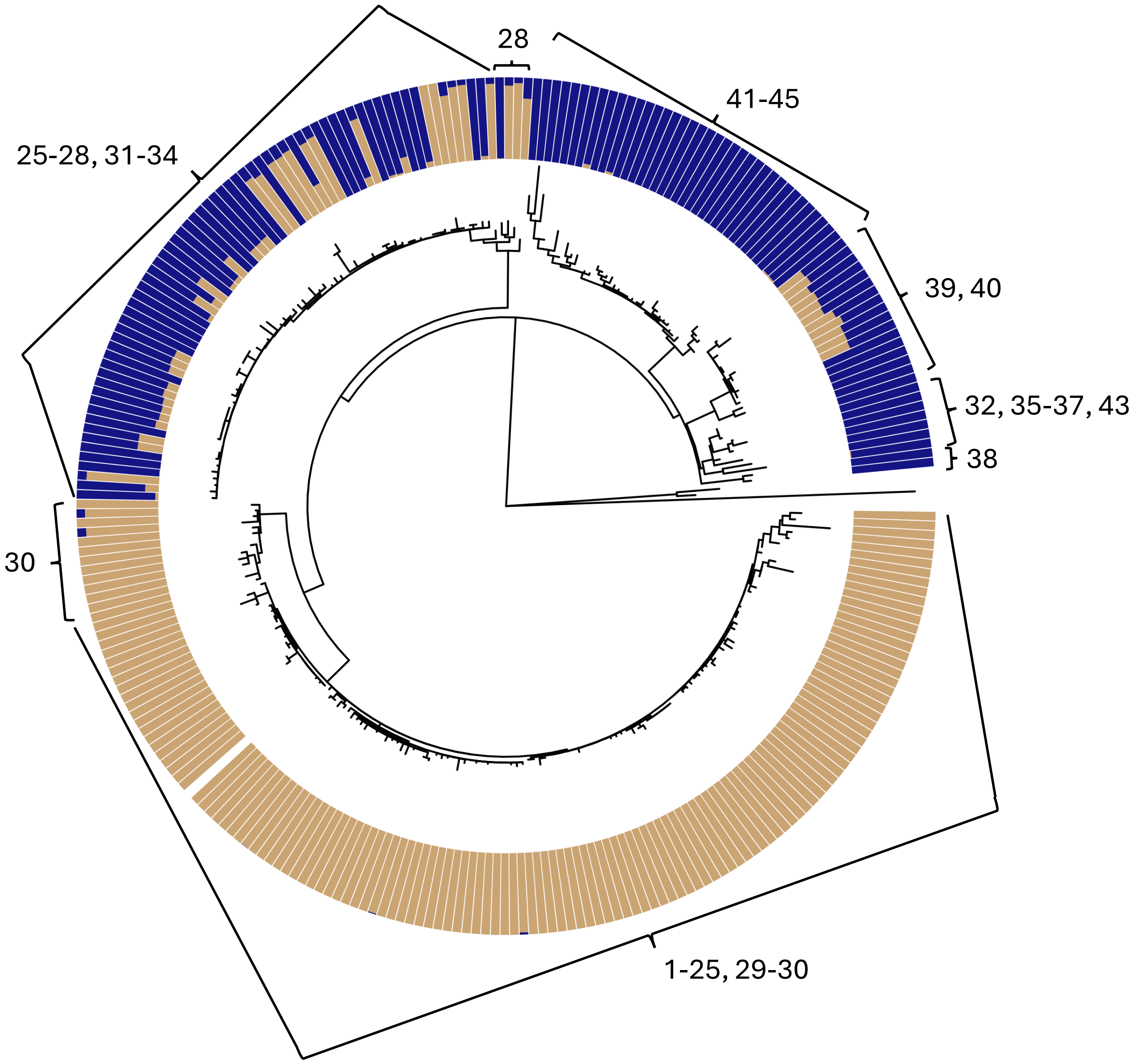


**Figure S2**: The internal circle shows a maximum likelihood tree of the mitochondrial genome of the *F. podzolica* samples in the whole genome sequencing dataset, with two *F. glacialis* individuals included as an outgroup. The circular barplot shows the nuclear genome-based results from Admixture (K = 2). The numerical annotations on the outside of the figure indicate the sampling localities of individuals from each major mitochondrial clade.

Supplementary Materials for: Purcell et al. Variation in social organization and supergene control along a latitudinal gradient

All tables are included in the accompanying spreadsheet file, which contains one tab for each table. The tables are as follows:

**Table S1**: This table contains a list of samples included in the whole genome sequencing dataset, including individual ID, caste, state or province collected, latitude, longitude, supergene genotype, mean read depth, and geographic locality. Additional technical information about each sample is provided in the NCBI SRA, project PRJNA1293783.

**Table S2**: This table contains a list of colonies included in the RADseq analysis, including the colony ID, the state or province collected, latitude, longitude, supergene genotypes found in the colony, number of individuals assessed, colony social structure (mono is monogyne and monandrous, mono2 is monogyne and polyandrous, and poly is polygyne), sequencing batch, and geographic locality.

**Table S3**: A list of the 200 genes in the first cassette, which spans the '$M_D$' region, extending from 2.05 to 5.37 Mbp along chromosome 3.

**Table S4:** A list of the 172 genes in the second cassette, which extends from 14.06 to 17.6 Mbp along chromosome 3.

**Table S5:** A list of the 115 genes in the third cassette, which contains the gene *Knockout* and extends from 17.6 to 19.4 Mbp along chromosome 3.

**Table S6:** A list of the 77 genes that are highly differentiated between the northern and southern *F. podzolica* populations.
